# Significance of two transmembrane ion gradients for human erythrocyte volume stabilization

**DOI:** 10.1101/2022.07.26.501528

**Authors:** F.I. Ataullakhanov, M.V. Martinov, Qiang Shi, V.M. Vitvitsky

## Abstract

Functional completeness of erythrocytes depends on high deformability of these cells, that allows them to pass through narrow tissue capillaries. The erythrocytes high deformability is provided due to maintenance of discoid shape with an optimal cell surface area to volume ratio. This ratio can be maintained due to cell volume stabilization at a given cell surface area. We studied role of Na/K-ATPase and transmembrane Na^+^ and K^+^ gradients in human erythrocyte volume stabilization at non-selective increase in cell membrane permeability to cations by using mathematical simulation. The simulation took into account a contribution of glycolytic metabolites and adenine nucleotides to cytoplasm osmotic pressure in the cells. It was shown that in the presence of Na/K-ATPase activated by intracellular sodium ions and two oppositely directed gradients of Na^+^ and K^+^ ions in the cell, the volume of the erythrocyte deviates from the optimal value by no more than 10% with a change in the non-selective permeability of the cell membrane to cations from 50 to 200% of the normal value. The transport Na/K-ATPase, which sets the ratio of transmembrane fluxes of sodium and potassium ions equal to 3:2, provides the best stabilization of the erythrocyte volume exactly at a non-selective increase in the permeability of the cell membrane, when the permeability for sodium and potassium ions increases equally. Such increase in erythrocyte membrane permeability is caused by oxidation of the membrane components and by mechanical stress during circulation. In the case of only one transmembrane ion gradient (Na^+^), the cell loses the ability to stabilize the volume when the cell membrane is damaged. In this case even small variations of cell membrane permeability cause dramatic changes in the cell volume. Our results reveal that the presence of two oppositely directed transmembrane ion gradients (Na^+^ and K^+^) and the transport Na/K-ATPase activated by intracellular sodium are fundamentally important conditions for the stabilization of cellular volume in human erythrocytes.

## Introduction

The main function of erythrocytes is oxygen transportation from lungs to tissues. This function is provided due to erythrocytes ability to bind big amounts of oxygen and to circulate in the bloodstream passing through narrow tissue capillaries. The ability of erythrocytes to reversibly bind a large amount of oxygen is determined by a high concentration of hemoglobin in these cells (about 300 g per liter of cells [1]) and does not require energy costs. However, the ability of mammalian erythrocytes to circulate in the bloodstream depends on their ability to pass through narrow tissue capillaries which diameter is smaller compared with the dimension of the erythrocytes [2–4]. And this ability is energy-dependent.

Mammalian erythrocytes passing through the narrow tissue capillaries easily deform, changing their shape [2]. Erythrocytes with reduced deformability are removed from the bloodstream mainly in the spleen [4–7]. Thus, high deformability is the main criterion which determines mammalian erythrocytes usefulness and viability in an organism. The high deformability of mammalian erythrocytes is provided due to the presence of an elastic, but practically inextensible cell membrane and the discoid shape of these cells [4,8]. Normal human erythrocytes have the shape of a biconcave disc with a diameter of 7-8 microns and a thickness of about 2 microns [9]. The discoid shape of an erythrocyte means that its cellular volume is significantly smaller than the volume of a sphere with a surface area equal to the surface area of its cell membrane. The volume of normal human erythrocytes is maintained within 55-60% of the maximum volume that a sphere with the same surface area as an erythrocyte has [10]. In other words, normally an erythrocyte has an excess surface relative to its volume. With an increase in cell volume by 1.7-1.8 times compared to normal, the erythrocyte takes the form of a sphere and, as a result, loses the ability to deform [3,4,10]. Since the erythrocyte membrane is inextensible, a further increase in cell volume leads to cell membrane rupture and destruction of the cell [4]. On the other hand, a decrease in the volume of the erythrocyte leads to an increase in the concentration of hemoglobin in the cytoplasm. As a result, the viscosity of the cytoplasm increases, and such erythrocyte also loses the ability to deform and to pass through narrow tissue capillaries [3,4,10,11]. Hard disk-shaped erythrocytes are unacceptable for a circulation as well as spherical cells. Therefore, the circulating erythrocytes should maintain an optimal ratio of the cell surface area to its volume. In the blood of a normal healthy donor, the cellular volume and surface area of circulating erythrocytes may vary more than two times, while the deviations of the ratio of surface area to volume for individual red blood cells lies within ± 5% of the average value [7,12–14]. In fact, this value is stabilized within the experimental error in all circulating erythrocytes. Since cells have a number of cell volume regulating systems [15], it is reasonable to assume that the erythrocyte stabilizes its volume at a given cell membrane area so as to obtain an optimal ratio of surface area to volume.

Human erythrocyte volume depends on osmotic pressure. The erythrocyte membrane permeability to water is very high [16], and the intracellular concentration of proteins and metabolites that do not penetrate through the cell membrane is significantly higher than in blood plasma. Rough estimates show that the total difference in the concentration of osmotically active components that do not penetrate the cell membrane in erythrocytes, compared with blood plasma, is about 50 mM (50 mOsm) [17,18]. This should lead to increased osmotic pressure inside the cells. Animal cells do not try to resist osmosis. They equalize the osmotic pressure on both sides of the cell membrane, because the cell membrane breaks easily when stretched and cannot hold a pressure exceeding 2 kPa (∼1 mOsm) [19]. In order to compensate for the osmotic pressure caused by macromolecules and metabolites, the cell could reduce the intracellular concentration of some other substances, of which there are quite a lot both inside and outside the cell. In most cells, sodium ions are used as such a substance. And it would be quite natural to have a pump that removes only sodium ions from the cell to decrease an intracellular sodium concentration and to compensate a passive sodium transport to cytoplasm from the medium. To equalize the osmotic pressure between the cell and the medium, it would be enough to reduce the sodium ions concentration in the cell by about 50 mM compared to the medium. In fact, the concentration of sodium ions in the cell is reduced significantly more. At the same time, the cell performs seemingly meaningless work, pumping potassium ions into the cell in almost the same amount. But actually, it makes sense and below we will try to demonstrate the importance of the existence of two oppositely directed ion gradients between the cell and the medium.

Thus, the volume of an erythrocyte is a dynamic variable and can change quite easily with a change in the distribution of ions between the cell and the medium. To maintain its volume, the erythrocyte must maintain ion homeostasis. In human erythrocytes, the necessary distribution of ions between the cytoplasm and the external medium is created by an ion pump – transport Na/K-ATPase, which transfers K^+^ ions into the cell, and Na^+^ ions from the cell to the medium in a ratio of 2:3 [20,21], thereby reducing the total content of monovalent cations in the cell compared to the medium. As a result, the osmotic pressure outside and inside the cell is equalized. It should be noted that in the stationary state, the active fluxes of Na^+^ and K^+^ through the cell membrane due to the operation of the transport Na/K-ATPase should be equal to the passive transmembrane leakage of these ions (Fig.1).

**Fig. 1.**
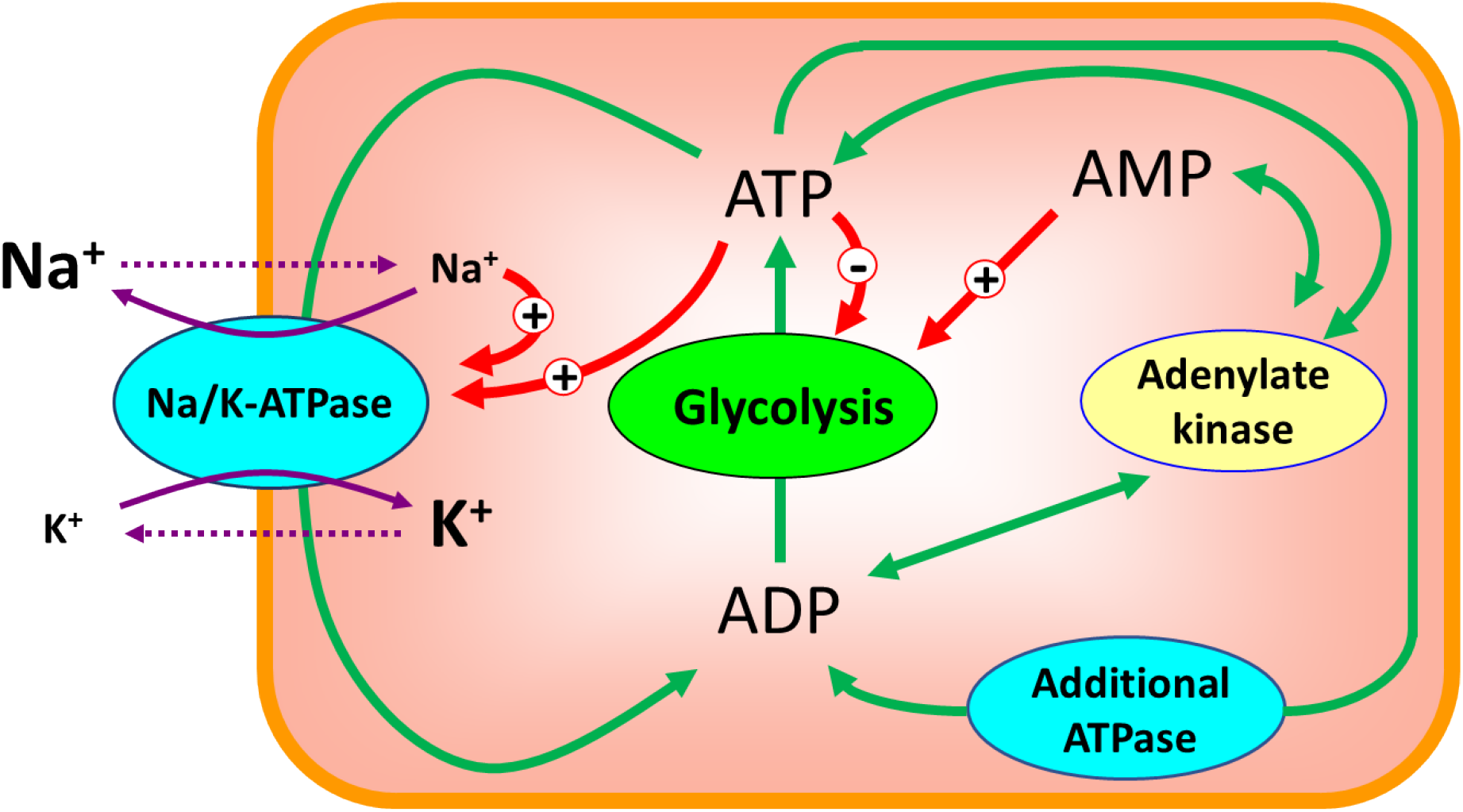
Interaction of transport Na/K-ATPase and glycolysis in human erythrocytes. Solid and dotted purple arrows show active and passive ion fluxes through the cell membrane respectively. Ion symbol size inside and outside the cell is proportional to the ion concentration. The red arrows show the activation (+) and inhibitory (-) effects of ions and adenylates on Na/K-ATPase and on glycolysis. The green arrows show the interconversions between ATP, ADP and AMP. The additional ATPase represents the ATP consuming processes in the cell other than the active transmembrane Na^+^ and K^+^ transport.

A number of transport systems capable of normalizing the volume of cells placed in a hypotonic or hypertonic environment have been described in the literature [15]. However, the composition of the extracellular medium in the body is well stabilized and a change in its osmotic activity is rather an exception than a common situation. That explains why erythrocyte is not protected against variations in osmolarity of an external medium and behaves in vitro as an ideal osmometer. The changes in the biochemical parameters of erythrocytes, such as Na/ K-ATPase activity, ATP concentration, etc., during their aging in the bloodstream (during circulation) occur slowly and values of the parameters, in principle, can ensure the maintenance of optimal cell volume throughout the lifetime of the erythrocyte. The question arises, what else can affect the volume of circulating red blood cells, or from what influences (disorders / damage) should these cells be primarily protected by the volume stabilization systems available in them?

One of the causes for a significant change in the erythrocyte volume in the body may be a disturbance of the permeability of the cell membrane for cations. During circulation, the erythrocyte membrane is exposed to high oxygen concentrations and undergoes to a significant mechanical stress, that can lead to a non-selective increase in its permeability to cations, that is, to the same increase in permeability for all cations [22–29]. Experimental and theoretical studies show that this may lead to an increase in cell volume and to the lysis of erythrocytes [17,25,26,30].

Despite the presence in erythrocytes of a number of systems capable of regulating cell volume [15], the possibility of participating in the stabilization of cell volume at a non-selective increase in the permeability of the cell membrane has not been demonstrated for any of them. Moreover, possible mechanisms of erythrocyte volume regulation are discussed in the literature mainly at the descriptive level. Earlier, using mathematical modeling, we showed that the transport Na/K-ATPase can provide stabilization of the erythrocytes cell volume at a non-selective increase in the permeability of the cell membrane for cations [17,31]. However, the mathematical models used did not take into account the contribution of glycolysis metabolites and adenine nucleotides to the osmotic pressure of the cytoplasm. Here, using the updated model, which takes into account the contribution of glycolysis metabolites and adenine nucleotides to the osmotic pressure of the cytoplasm, we have shown that the transport Na/K-ATPase provides the best stabilization of the erythrocyte volume exactly at a non-selective increase in the permeability of the cell membrane. Moreover, we have shown that the presence of two oppositely directed transmembrane ion gradients (Na^+^ and K^+^) is crucial for a good sensitivity of the cell to the cell membrane damage and for stabilization of the cell volume at non-selective variations of the cell membrane permeability to cations.

## Results

Mathematical modeling of the human erythrocytes volume regulation has shown that in the presence of transport Na/K-ATPase in the cell, a non-selective increase in the cell membrane permeability leads to an increase in cell volume due to an increase in the intracellular concentration of Na^+^ (Fig. 2 A, B). The increase in intracellular [Na^+^] is compensated, in part, by a decrease in the intracellular [K^+^]. Also, the increase in the intracellular sodium concentration leads to an activation of the transport Na/K-ATPase (Fig. 3A), which, in turn, leads to a compensation of the increased passive transmembrane fluxes of Na^+^ and K^+^ caused by an increase in the permeability of the cell membrane. As a result, with a twofold increase in membrane permeability, the volume of the erythrocyte increases by only 10% compared to the initial value (Fig. 2A). When cell membrane permeability increases by 5 times, the volume of the erythrocyte reaches the maximum value corresponding to the spherical shape of the cell (Fig. 2A), at which it completely loses the ability to deform and, consequently, to circulate in the bloodstream. Any further increase in the erythrocyte volume causes its disruption. The results presented here are in good agreement with the results obtained earlier using models that do not take into account the contribution of glycolysis metabolites and adenylates to the osmotic pressure of the cytoplasm [17,31]. Thus, in our conditions, glycolysis metabolites and adenylates do not significantly affect the regulation of cell volume in erythrocytes.

**Fig. 2.**
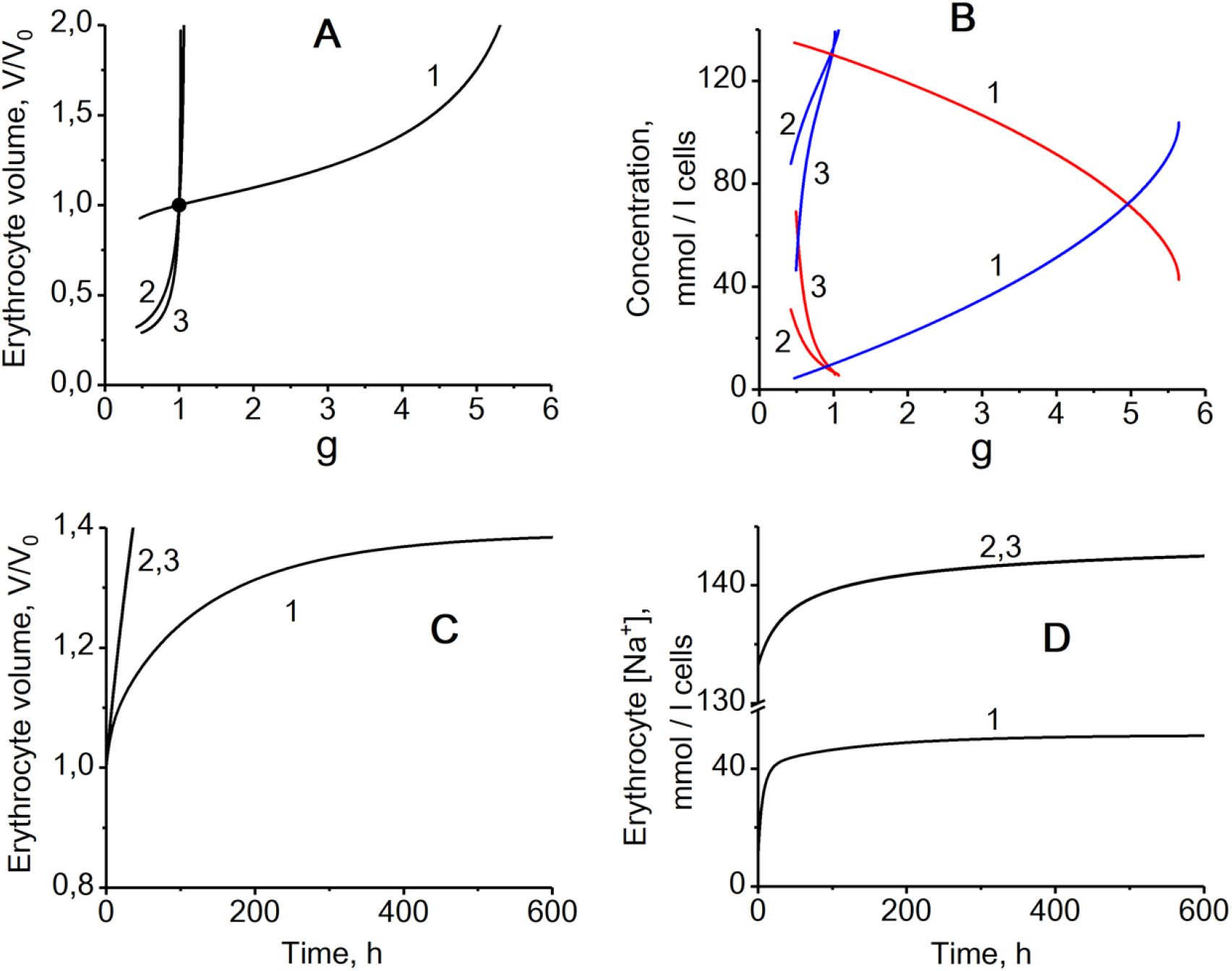
The effect of nonselective permeability of the cell membrane for cations on the erythrocyte volume and on intracellular Na^+^ and K^+^ concentrations in different models. The dependence of the relative stationary volume of the erythrocyte (A) and stationary intracellular Na^+^ (blue lines) and K^+^ (red lines) concentrations (B) on the relative nonselective permeability of the cell membrane for cations 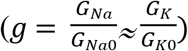. Kinetics of changes in erythrocyte volume (C) and intracellular Na^+^ concentration (D) after an instant 4-fold increase in the nonselective permeability of the cell membrane for cations. The kinetics of the K^+^ concentration is the same as for Na^+^, but changes occur in the direction of decreasing concentration. The black circle in the panel A indicates physiologically normal state of erythrocyte. The numbers on the curves correspond to the model versions: - 1 – Version 1, the basic version of the model with actively maintained transmembrane Na^+^ and K^+^ gradients and transport Na/K-ATPase activated by intracellular sodium ions; 2 – Version 2, with actively maintained transmembrane gradient only for Na^+^ and sodium-activated transport Na-ATPase activated by intracellular sodium ions; 3 – Version 3, with actively maintained transmembrane gradient only for Na^+^ and transport Na-ATPase independent on intracellular sodium ions.

**Fig. 3.**
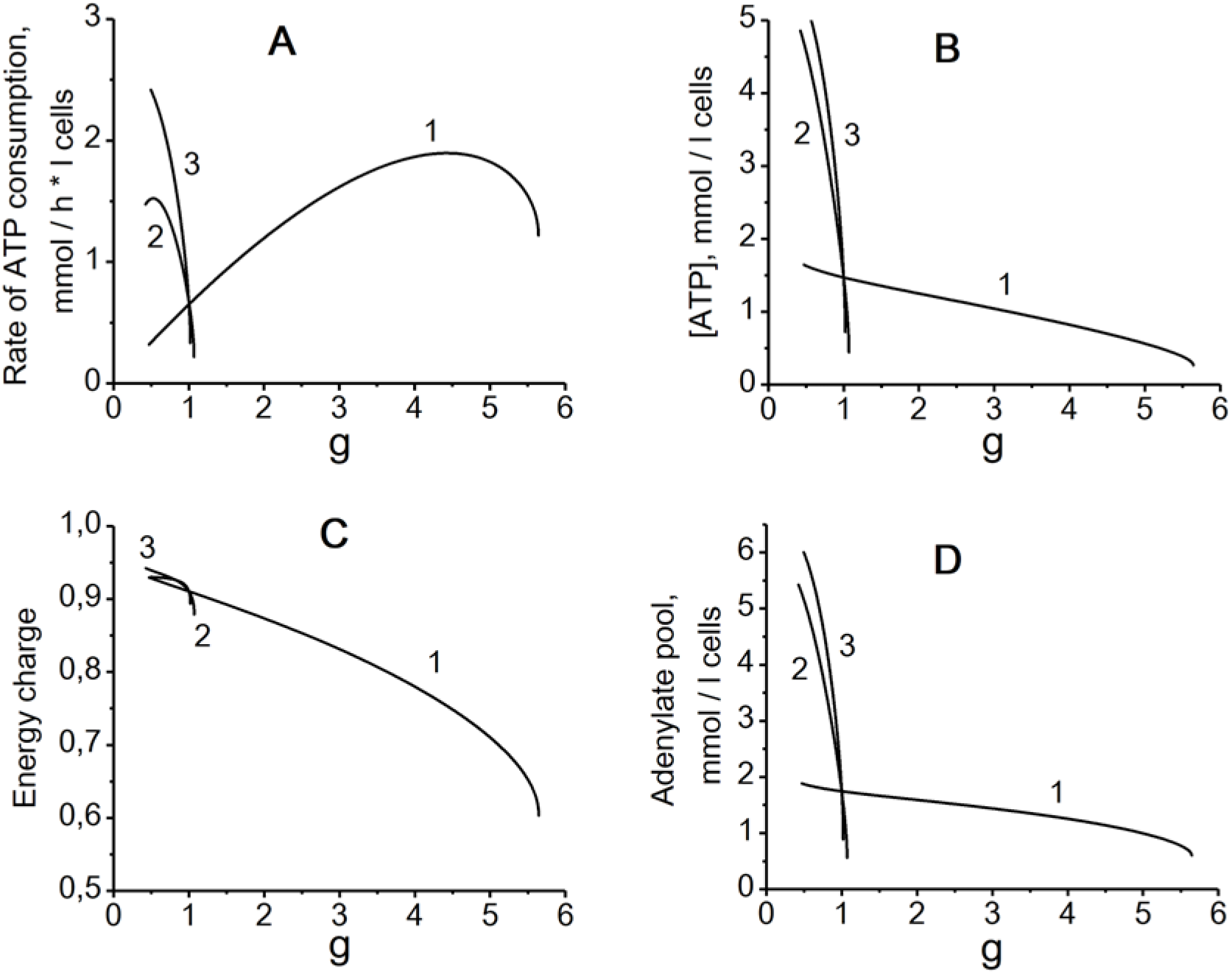
The effect of nonselective permeability of the cell membrane on erythrocyte energy metabolism in different models. (A) - The steady-state rate of ATP consumption by ion pumps; (B) - ATP concentration; (C) – Energy charge (([ATP] + 0.5[ADP])/([ATP] + [ADP] + [AMP])); (D) – Adenylate pool ([ATP] + [ADP] + [AMP]). The numbers on the curves indicate the model versions in the same way as in Fig. 2.

Now let us consider a model which includes only one active transmembrane ion gradient (Na^+^) and an ion pump which transports only sodium ions from the cell to a medium (Na-ATPase). The rate of this ATPase is proportional to concentrations of sodium ions and ATP (Equation 21). Here we assume a passive distribution of potassium ions between cytoplasm and blood plasma in accordance with transmembrane potential. In this case normal steady-state intracellular [K^+^] is low and close to the extracellular one (Fig. 2B). In a such model, a stationary value of the cell volume is also established, but actually there is no stabilization of this volume at a non-selective change in the permeability of the cell membrane for cations (Fig. 2A). Even small variations of cell membrane permeability cause dramatic changes in the cell volume and intracellular ion concentrations (Fig. 2A, B). Indeed, the difference in the concentration of osmotically active components between the erythrocyte and the medium is about 50 mM, that is, about 17% of the total concentration of osmotically active components in the cell [17,18] (Table 1). Thus, in the case of a single active transmembrane ion gradient, this gradient should be relatively small. The extracellular sodium concentration is about 150 mM and it should be just about 50 mM lower inside the cell. And with a small gradient, it is impossible to achieve a significant increase in the concentration of Na^+^ ions in the cell when the cell membrane is damaged (Fig. 2B), that is necessary for the effective activation of the transport Na-ATPase. Actually, the rate of transport Na-ATPase in the model decreases with an increase in cell membrane permeability due to a decrease in ATP concentration caused by an increase in cell volume and dilution of adenylates (Fig. 3). Of course, even worse cell volume stabilization was obtained in the model which includes only one transmembrane ion gradient (Na^+^) and a transport sodium pump (Na-ATPase) which depends on [ATP] but is independent on sodium concentration (Fig. 2, 3).

**Table 1.**
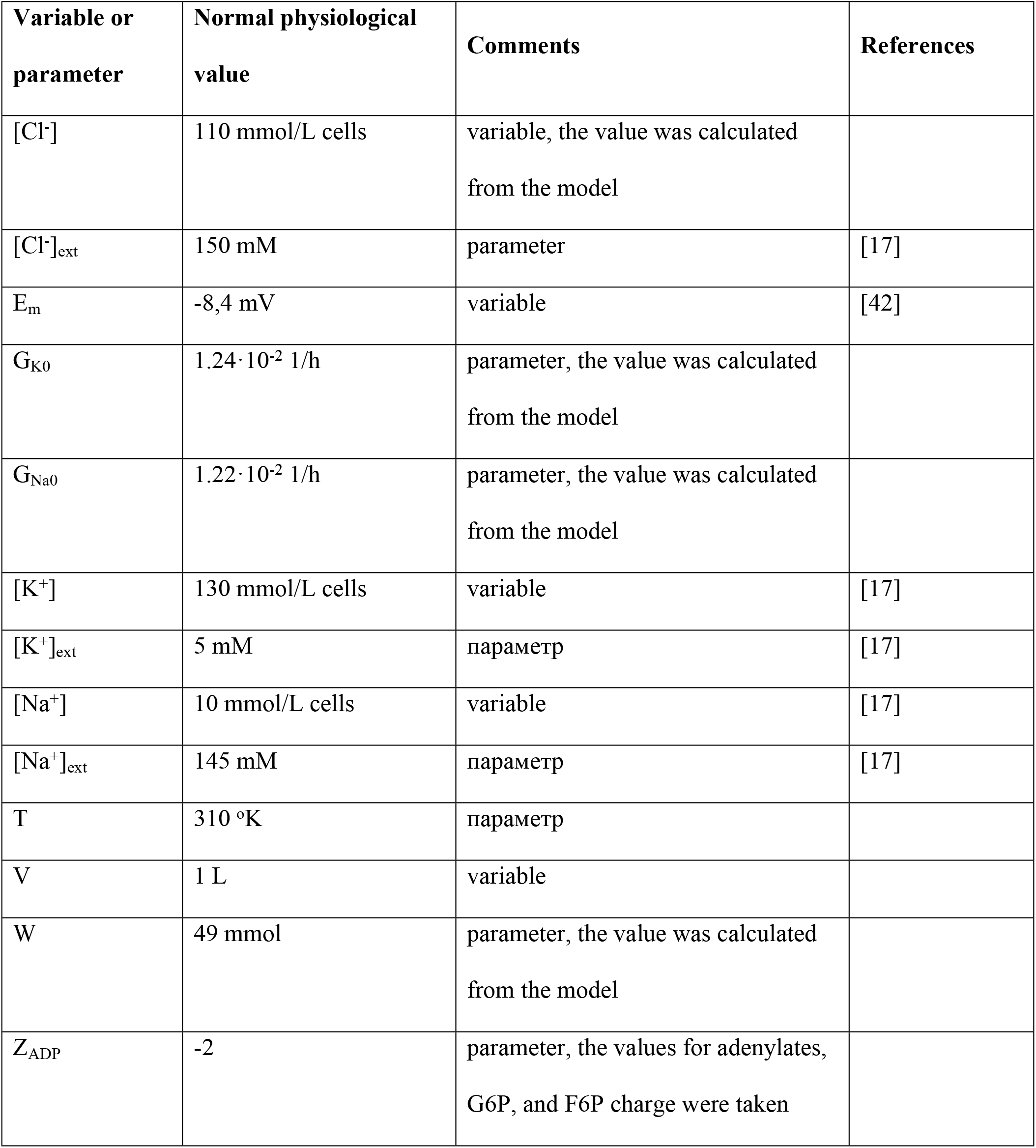
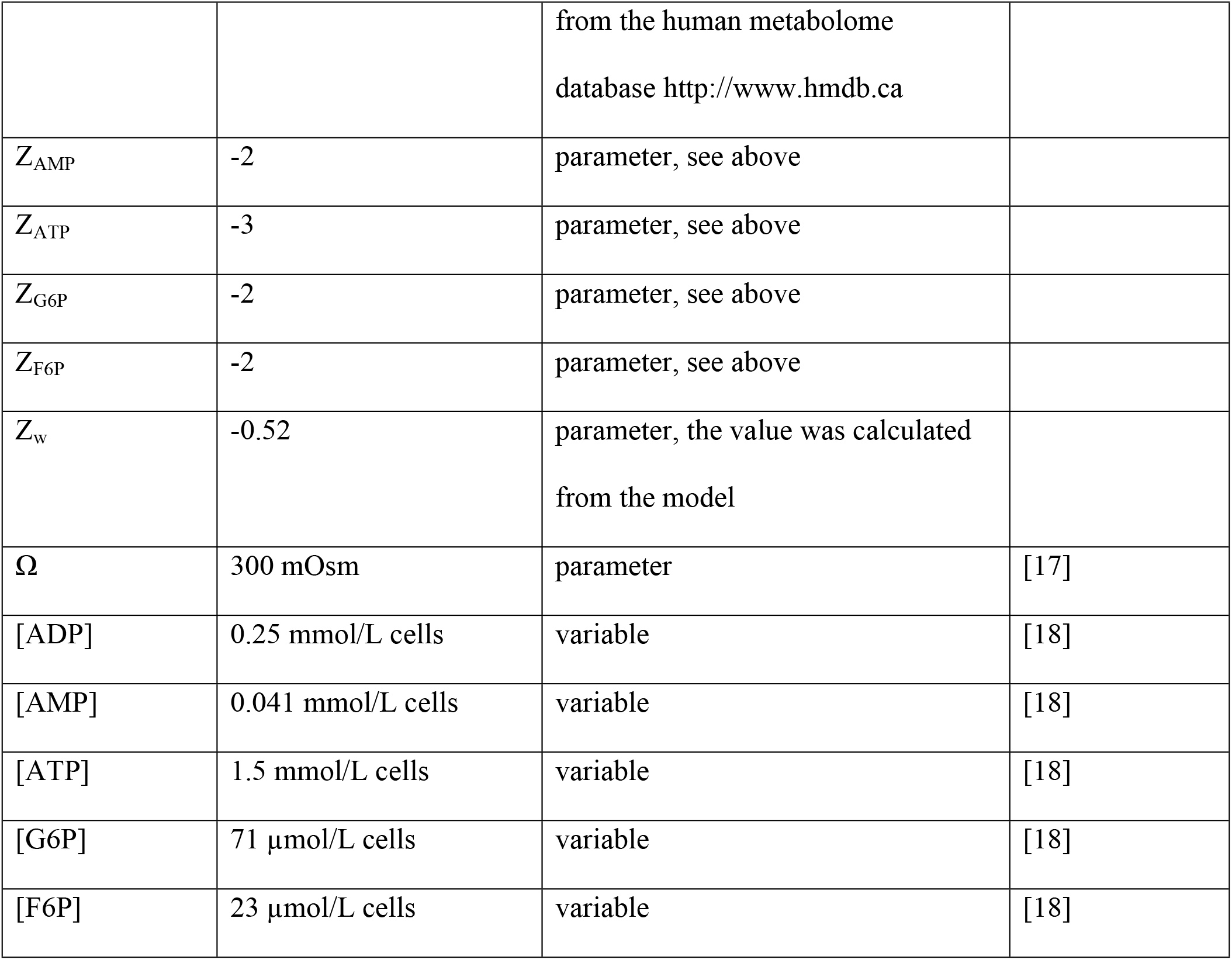
Model variables and parameters characterizing the normal physiological state of a human erythrocyte.

| Variable or parameter | Normal physiological value | Comments | References |
| --- | --- | --- | --- |
| $[Cl^-]$ | 110 mmol/L cells | variable, the value was calculated from the model | |
| $[Cl^-]_{ext}$ | 150 mM | parameter | [17] |
| $E_m$ | -8,4 mV | variable | [42] |
| $G_{K0}$ | $1.24 \cdot 10^{-2}$ 1/h | parameter, the value was calculated from the model | |
| $G_{Na0}$ | $1.22 \cdot 10^{-2}$ 1/h | parameter, the value was calculated from the model | |
| $[K^+]$ | 130 mmol/L cells | variable | [17] |
| $[K^+]_{ext}$ | 5 mM | параметр | [17] |
| $[Na^+]$ | 10 mmol/L cells | variable | [17] |
| $[Na^+]_{ext}$ | 145 mM | параметр | [17] |
| T | 310 °K | параметр |  |
| V | 1 L | variable |  |
| W | 49 mmol | parameter, the value was calculated from the model |  |
| $Z_{ADP}$ | -2 | parameter, the values for adenylates, G6P, and F6P charge were taken from the human metabolome<br>database http://www.hmdb.ca | |
| $Z_{AMP}$ | -2 | parameter, see above | |
| $Z_{ATP}$ | -3 | parameter, see above | |
| $Z_{G6P}$ | -2 | parameter, see above | |
| $Z_{F6P}$ | -2 | parameter, see above | |
| $Z_w$ | -0.52 | parameter, the value was calculated<br>from the model | |
| $\Omega$ | 300 mOsm | parameter | [17] |
| [ADP] | 0.25 mmol/L cells | variable | [18] |
| [AMP] | 0.041 mmol/L cells | variable | [18] |
| [ATP] | 1.5 mmol/L cells | variable | [18] |
| [G6P] | 71 μmol/L cells | variable | [18] |
| [F6P] | 23 μmol/L cells | variable | [18] |

In the case of two opposite gradients of Na^+^ and K^+^ (that is the case in most mammalian cells), the difference in the sum concentration of Na^+^ and K^+^ ions between the cell and the medium is the same 50 mM, but the intracellular concentration of Na^+^ is many times less than the concentration of Na^+^ in the medium. This leads to the fact that when the cell membrane is damaged, the concentration of Na^+^ ions in the cell can significantly exceed the physiologically normal value (Fig. 2B) and, consequently, cause significant activation of the transport Na/K-ATPase (Fig. 3A) that provides cell volume stabilization. Thus, the presence of opposite gradients of Na^+^ and K^+^ between the cytoplasm and the medium allows the cell to respond effectively to a damage of the cell membrane and stabilize the cell volume by activating the transport Na/K-ATPase. It is the large transmembrane gradient of sodium ions that ensures the rapid and significant increase in its concentration in the cytoplasm when the cell membrane is damaged. And this gradient is achieved due to the presence of an oppositely directed transmembrane gradient of potassium ions.

We also found that the transport Na/K-ATPase, which sets the ratio of transmembrane fluxes of sodium and potassium ions equal to 3:2, provides the best stabilization of the erythrocyte volume exactly at a non-selective increase in the permeability of the cell membrane, when the permeability for sodium and potassium ions increases equally (Fig. 4). The cell volume stabilization getting significantly worse if the cell membrane permeability increases predominantly for one of the ions. As one can see from Fig. 4, the erythrocyte volume increases significantly if the permeability of the cell membrane for Na^+^ increases at constant permeability for K^+^. Contrary, the erythrocyte volume decreases significantly if the membrane permeability for K^+^ increases while the permeability for Na^+^ is constant. If both membrane permeabilities increase simultaneously the cell volume almost does not change. Thus, the cell is best protected from a non-selective increase in the cell membrane permeability.

**Fig. 4.**
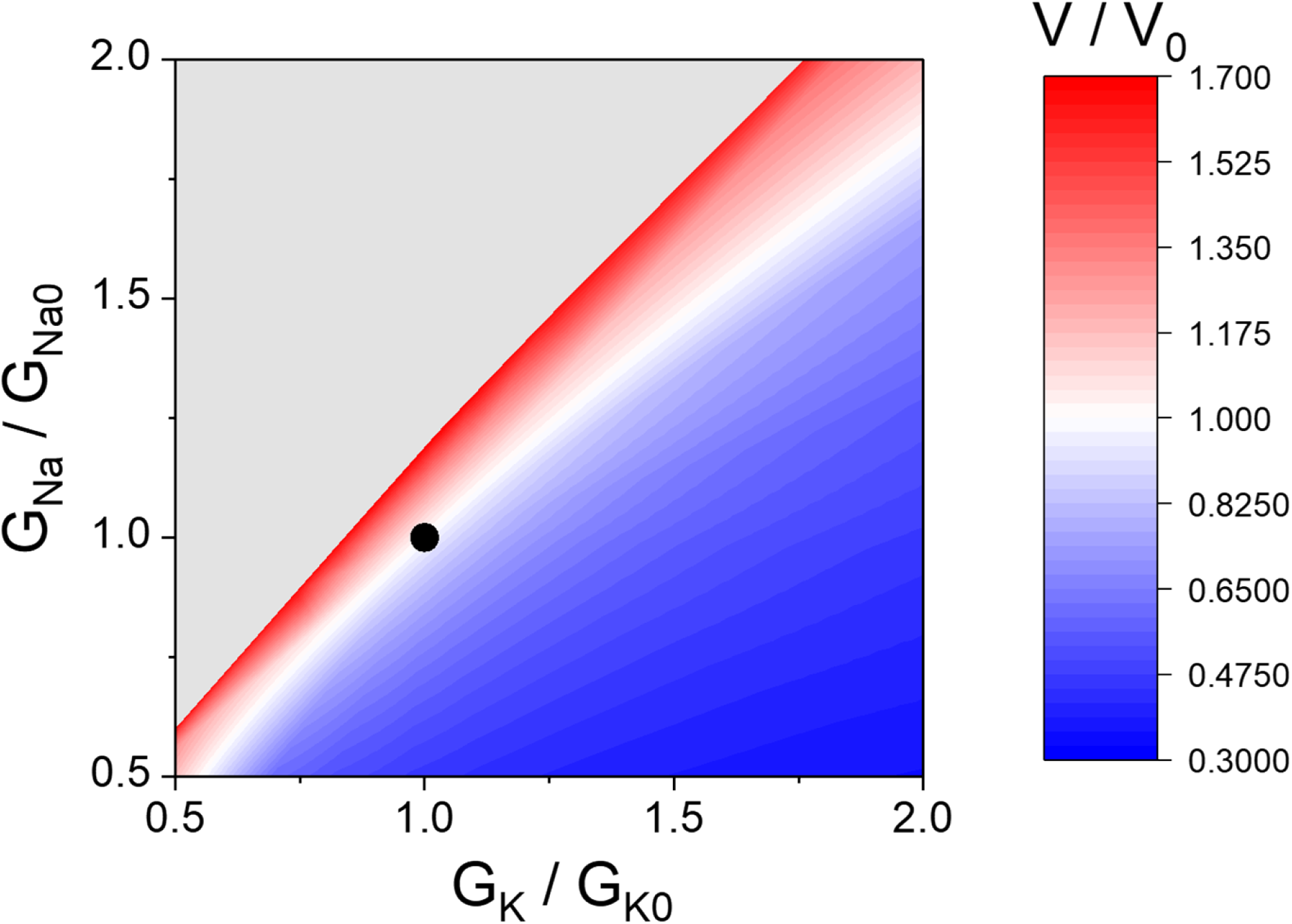
The dependence of the relative steady-state erythrocyte volume (V/V_0_) in the model on the passive permeability of the cell membrane for potassium (G_K_/G_K0_) and sodium (G_Na_/G_Na0_) ions. The model includes actively maintained transmembrane Na^+^ and K^+^ gradients and transport Na/K-ATPase activated by intracellular sodium ions (Version 1 - the basic version of the model). A black circle indicates the normal physiological state of the erythrocyte.

Interestingly, in the case of only one transmembrane ion gradient, a non-selective decrease in the permeability of the cell membrane leads to a strong decrease in cell volume (сurves 2 and 3 in Fig. 2A). In all models, a change in the non-selective permeability of the cell membrane leads to a more or less significant change in the cell volume (Fig. 2A). In turn, this leads to changes in the adenylate pool value, although the amount of adenylates in the cell remains constant in the models (Fig. 3D).

The characteristic time for establishing a new steady state in the model after changing the permeability of the cell membrane is tens or even hundreds of hours and is determined by the low total rate of ion fluxes through the cell membrane (Fig. 2C, D). Moreover, in the case of two transmembrane ion gradients, the release of potassium ions from the cells partially compensates for the entry of sodium ions into the cells and also slows down the rate of change in cell volume.

It should be noted that the stabilization of the erythrocyte volume only due to the transport Na/K-ATPase is associated with significant changes in intracellular levels of Na^+^ and K^+^ (Fig. 2B), that is, with a disturbance of ion homeostasis in the cell, as well as with changes in the levels of ATP and the energy charge (([ATP] + 0.5[ADP])/([ATP] + [ADP] + [AMP])) in the cell (Fig. 3B, C).

## Discussion

Our results show that the presence of two oppositely directed active transmembrane ion gradients (Na^+^ and K^+^) and the transport Na/K-ATPase activated by intracellular sodium are fundamentally important conditions for the stabilization of cellular volume in human erythrocytes. Under these conditions, the most effective stabilization of the cell volume is provided at a non-selective increase in the permeability of the cell membrane. It seems to us that a non-selective increase in the permeability of erythrocyte membranes is the most likely damage to the cell membrane of these cells under conditions of circulation in the bloodstream. Based on these results, a general conclusion can be done that the presence of two oppositely directed transmembrane ion gradients (Na^+^ and K^+^) at a low intracellular concentration of the ion prevailing in the external medium (Na^+^) provides a greater (compared to conditions with a single gradient (Na^+^)) sensitivity of the cell to a damage of the cell membrane and is a fundamentally necessary condition for ensuring the cell volume stabilization. One can assume that such cell organization arose in the early stages of evolution and later served as the basis for the emergence of cellular electrical excitability, etc. Also, this result casts doubt on the hypothesis that the high level of potassium in cells reflects the ionic composition of the environment in which the first cellular organisms originated [32–34]. From the results presented here, it follows that in order to preserve the integrity of cells and maintain cellular volume, it would be very impractical for primary organisms to maintain an intracellular ionic composition similar to the composition of the external environment.

Nevertheless, Na/K-ATPase alone cannot provide stabilization of the erythrocyte volume in a sufficiently wide range of changes in the permeability of the cell membrane. Mathematical simulation shows that at two times increase in membrane permeability compared to the normal value, the cell volume increases by only 10% (Fig. 2A). However, this already goes beyond the ±5% frame in which the volume of the erythrocyte to its surface area ratio is stabilized in the body [7,12–14]. An increase in the permeability of the cell membrane by more than 5 times leads to the destruction of the erythrocyte. At the same time, some literature data indicate that erythrocytes can remain in the bloodstream with an increase in the permeability of the cell membrane by more than 5-10 times compared to the normal value [35–38]. The results of our previous studies show that taking into account the additional ion transport system (calcium-activated potassium channels or Gardos channels) and the metabolism of adenine nucleotides in the model makes it possible to achieve almost perfect stabilization of the erythrocyte volume with an increase in the permeability of the cell membrane up to 15 times compared to the normal value [17,31]. In this regard, the role of Gardos channels and adenylate metabolism in stabilizing the volume of human erythrocytes should be revised using more correct models.

## Methods (Mathematical model description)

Mathematical model used in this study is a system of algebraic and ordinary differential equations which describe transmembrane ion fluxes, osmotic regulation of human erythrocyte cell volume, and glycolysis.

### Ions and Cell volume Regulation

The description of ion balance and cell volume regulation here is based on the mathematical model of human erythrocyte volume regulation published earlier in [17]. Equation (1) describes equivalence of osmolality in cytoplasm and in external medium (blood plasma) while the equation (2) describes the cytoplasm electroneutrality:

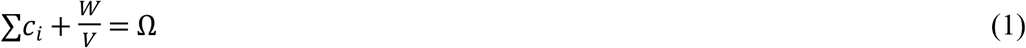

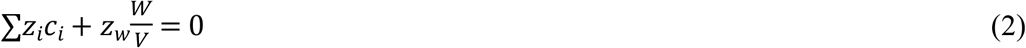

Here c_i_ denotes concentration of each osmotically active cytoplasm component, which is described in the model as a variable. They are Na^+^, K^+^, and Cl^-^ ions, adenylates – ATP, ADP, AMP, and glycolysis intermediates – glucose-6-phosphate (G6P) and fructose-6-phosphate (F6P). W denotes total amount of all other osmotically active components of human erythrocytes, including hemoglobin, enzymes, and metabolites which kinetics is not described explicitly in the model. Z_i_ denotes electrical charge of the corresponding osmotically active component. Z_W_ denotes the average electrical charge of components not explicitly described in the model. Ω - total concentration of osmotically active components in blood plasma. V – erythrocyte volume. It is more convenient to express volume in the model as per 10^13^ erythrocytes that is equal to one liter rather than per one erythrocyte. Thus, under normal physiological conditions V=V_0_=1 L. In this case, the amounts of substances expressed in grams in the cells are numerically equal to their molar concentrations.

Equations (3, 4) describe kinetics of quantity of Na^+^ and K^+^ ions in erythrocytes:

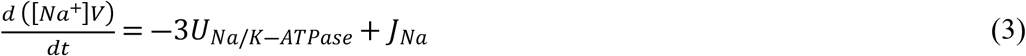

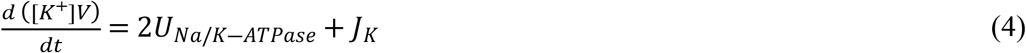

Here *U*_Na/K-ATPase_ – the rate of ATP consumption by transport Na/K-ATPase, *J*_*Na*_ and *J*_*K*_ are passive Na^+^ and K^+^ ion fluxes across cell membrane respectively. The cell membrane permeability for Cl^-^ anions in the erythrocyte is two orders of magnitude higher than for cations [30], therefore equation (5) describes the equilibrium distribution of chlorine between the cytoplasm and the medium in accordance with the electric potential on the membrane:

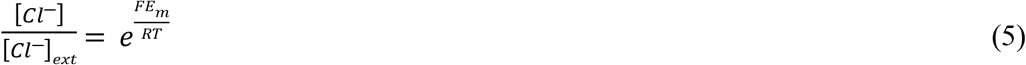

Here [Cl^-^] and [Cl^-^]_ext_ – intracellular and extracellular concentrations of chlorine ions respectively, F – Faraday constant, E_m_ - electrical potential on the erythrocyte cell membrane, R – universal gas constant, T - absolute temperature (310°K). Equation for the rate of Na/K-ATPase and parameter value were taken from [17]:

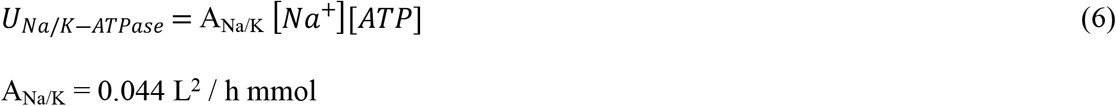

The passive ion fluxes across the erythrocyte cell membrane are described in Goldman approximation regarding constancy of the electric field inside the cell membrane [39]:

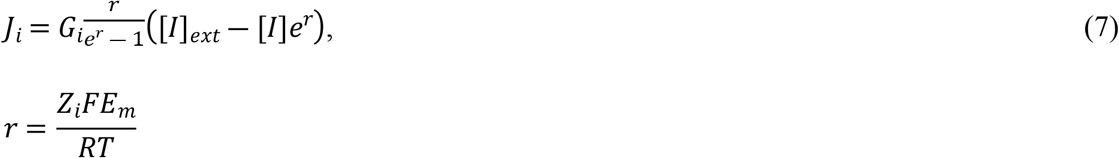

Here G_i_ – erythrocyte membrane permeability for ion I. [I] and [I]_ext_ – concentration of ion I inside the erythrocyte and in the external medium respectively.

### Glycolysis and energy metabolism

In the model we used the simplified description of glycolysis. Only the upper part of glycolysis is explicitly described, including hexokinase, glucosephosphateisomerase, and phosphofructokinase reactions - which determine the rate of the entire glycolysis. Equations (8, 9) describe kinetics of quantity of G6P and F6P molecules in the erythrocyte respectively:

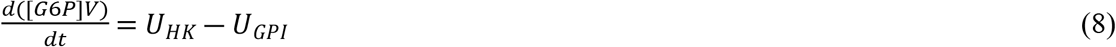

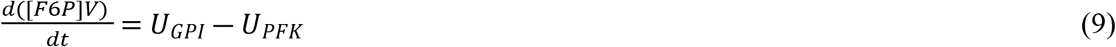

Here U_HK_, U_GPI_, and U_PFK_ denote hexokinase, glucosephosphateisomerase, and phosphofructokinase reaction rates in glycolysis respectively. We neglect the metabolic flow in the 2,3-diphosphoglycerate shunt and assume that the rate of reactions of the lower part of glycolysis (from aldolase to lactatedehydrogenase), is equal to twice the rate of the phosphofructokinase reaction. Then the rate of ATP production in glycolysis is determined by the difference between the rates of its production in phosphoglyceratekinase (U_PGK_) and pyruvate kinase (U_PK_) reactions and consumption in hexokinase and phosphofructokinase reactions:

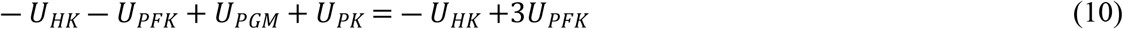

Total amount of ATP + ADP + AMP in the cells was assumed to be constant. We also assumed that rapid adenylate equilibrium exists in the cells all the time (Equation (11)).

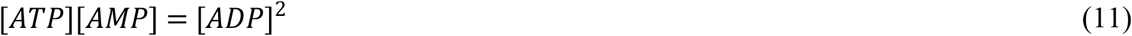

The energy balance in the cell can be described by the following equation:

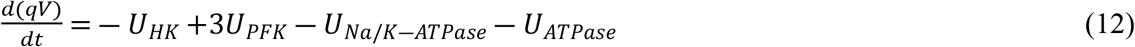

Here *q=* 2[ATP] + [ADP], U_ATPase_ – the rate of additional ATPase reaction added to the model to balance the rates of ATP production and consumption [17,18,40,41].

The presence of rapid adenylatekinase equilibrium in the cell leads to the fact that of the three variables – ATP, ADP, and AMP – only two are independent. The transition to the variables *q* and *p* = [ATP] + [ADP] + [AMP] (adenylate pool) makes it possible to exclude the rapid adenylate kinase reaction from the equations. Concentrations of ATP, ADP, and AMP at fixed values of *q* and *p* can be calculated from the following system of equations:

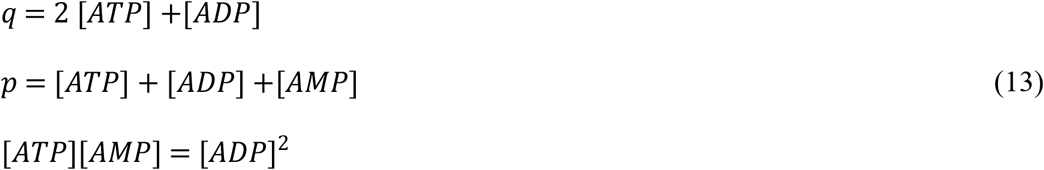

The equations for the rates of HK, GPI, PFK, and additional ATPase reactions and parameter values were taken from [18]:

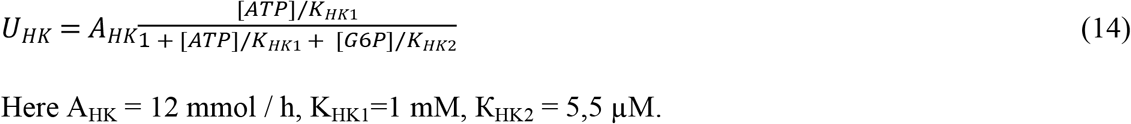

Here A_HK_ = 12 mmol / h, K_HK1_=1 mM, К_HK2_ = 5,5 µM.

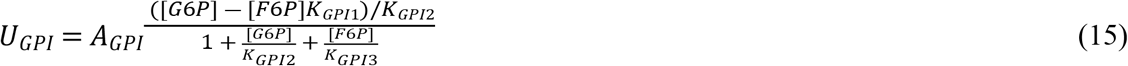

Here A_GPI_ = 360 mmol / h, К_GPI1_ = 3 mM, К_GPI2_ = 0.3 mM, К_GPI3_ = 0.2 mM

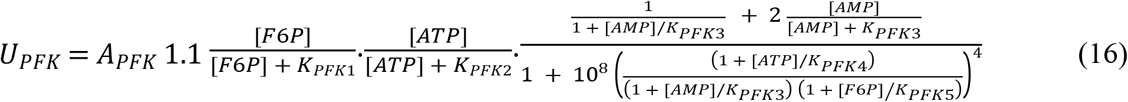

Here A_PFK_ = 380 mmol / h, К_PFK1_ = 0.1 mM, К_PFK2_ = 2 mM, К_PFK3_ = 0.01 mM, К_PFK4_ = 0.195 mM, К_PFK5_ = 0.37 µM.

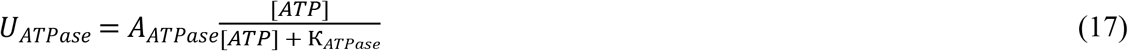

Here A = 2.7 mmol / h, K_ATPase_ = 1 mM

### Calculation of the model parameter values

To finalize the model, it is necessary to calculate some of its parameters – these are the normal physiological values of the intracellular concentration of chlorine ions, the permeability of the erythrocyte membrane for cations, the W value and the average charge of osmotically active components that do not penetrate the erythrocyte membrane (Z_W_), such as hemoglobin, enzymes and metabolites. First, we calculate the concentration of chlorine ions in the erythrocyte using equation (5) and taking the extracellular concentration of chlorine equal to 150 mM, the membrane potential equal to −8.4 mV and the temperature equal to 310 °K [17,42] (Table 1). Then, using equations (1) and (2), we calculate the values of W and Z_W_ using the physiological values of the variables from Table 1. According to these calculations W = 49 mmol and Z_W_ = 0.52.

### Reduction of a mathematical model to a system of ordinary differential equations

Initially the mathematical model includes five differential (3, 4, 8, 9, 12) and three algebraic (1, 2, 5) equations. If one chooses the amounts of substances in the erythrocyte as the model variables:

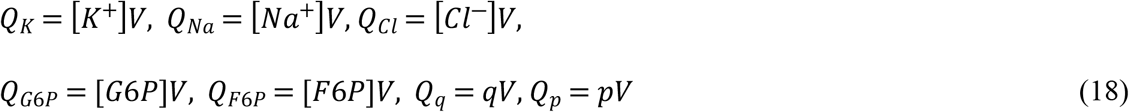

and express the variables V, Q_Cl_, and E_m_ through the remaining variables using algebraic equations (1, 2, 5), then the model is transformed into a system of five ordinary differential equations, similar to (3, 4, 8, 9, 12), in respect to variables Q_K_, Q_Na_, Q_G6P_, Q_F6P_, Q_q_.

Investigation of the kinetic behavior of the model was performed using the CVODE software [43], and investigation of the dependence of the steady state of the model on the parameters was performed using the AUTO software[44].

### Description of the cell membrane damage in the model

The main task of this work was to study the model in case of nonspecific damage of the cell membrane. As it is noted in the introduction, we assume that such damage leads to a nonselective increase in the permeability of the cell membrane for cations. In other words, the cell membrane permeability increases by the same amount for all cations:

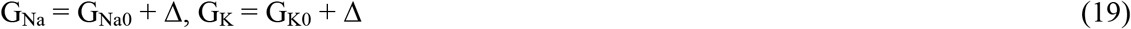

Here G_Na0_ and G_K0_ denote normal physiological values of cell membrane permeability for sodium and potassium cations respectively and Δ denotes a value of an increase in the membrane permeability for cations. Generally speaking, both an increase and a decrease in the cell membrane permeability can be described in this way. For the convenience of the results presentation, we introduced the parameter g corresponding to the relative permeability of the membrane for cations:

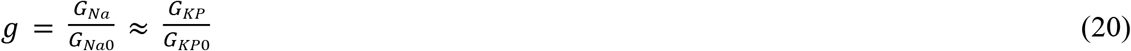

The validity of equation (20) follows from the fact that G_Na0_ ≈ G_K0_ (Table 1).

### The mathematical model versions

#### Initial model (Version 1)

The basic version of the model describing erythrocyte with transmembrane transport Na/K-ATPase, glycolysis, and constant adenylate content. The model consists of equations (3, 4, 8, 9, 12) modified according to the equations (18).

#### Version 2

The modified model (Version 1) in which the regular Na/K-ATPase was replaced with the transmembrane transport Na-ATPase (sodium pump which transports only sodium ions from the cell to external medium) which rate is proportional to intracellular concentrations of ATP and sodium ions. The rate of this pump is described by the following equation:

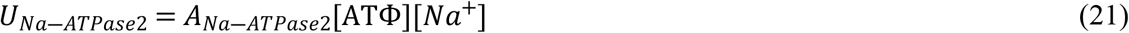

Here A_Na-ATPase2_ = 0.0035 L^2^ / h

The value of this parameter was chosen in such a way that this ATPase compensates for the passive flow of Na^+^ ions into the cell under physiological conditions.

In this version of the model the equation (4) does not contain a term describing active potassium transmembrane transport. Thus, the rate of change in intracellular potassium concentration is determined only by a passive potassium flux across the cell membrane according to the equation (7). Under these conditions the steady-state potassium concentration in the cells was calculated under assumption of zero passive potassium flux across the cell membrane according to the equation (7):

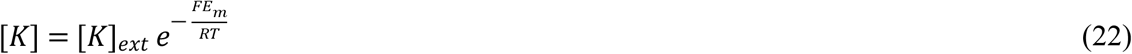

And the steady-state sodium concentration was calculated as follows:

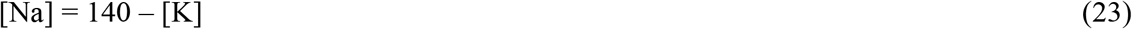

Here 140 (mmol/L cells) is a sum of sodium and potassium cation concentrations in erythrocytes containing Na/K-ATPase under normal conditions. This condition guarantees that the erythrocyte will have the same physiological volume and transmembrane potential in all model versions.

#### Version 3

The modified model (Version 1) in which the regular Na/K-ATPase was replaced with the transmembrane transport Na-ATPase (sodium pump which transports only sodium ions from the cell to external medium) which rate depends on [ATP] but does not depend on sodium concentration. The rate of this pump is described by the following equation:

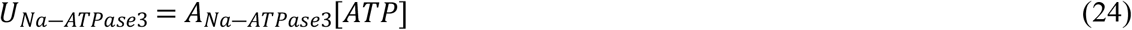

Here A_Na-ATPase3_ = 0.46 L / h

The value of this parameter was chosen in such a way that this ATPase compensates for the passive flow of Na^+^ ions into the cell under physiological conditions.

In this version of the model the intracellular potassium and sodium concentrations were calculated using the equations (23) and (24) the same as in Version 2.

## Author Contributions

F.I.A. general idea of the work and general supervision, analysis of literary data, construction of the mathematical models, and discussion of the results, M.V.M. all computer calculations, analysis of literary data, construction of the mathematical models, and discussion of the results, Q.S. discussion of the results, V.M.V. analysis of literary data, construction of the mathematical models, discussion of the results, writing the manuscript and preparing it for publication All the authors took part in the editing of the manuscript and approved its final version.

## Acknowledgments

This work was supported in part by grant from the Russian Science Foundation № 21-45-00012 (F.I. Ataullakhanov).

